# miRNA-mRNA network analysis identifies PAX5 as a potential regulator of adaptive immune response in COPD

**DOI:** 10.1101/2025.05.06.651484

**Authors:** Michele Gentili, Margherita De Marzio, Brian Hobbs, Craig P. Hersh, Min Hyung Ryu, Marieke L. Kuijjer, Michael H. Cho, Matthew Moll, Courtney Tern, Peter Castaldi, Kimberly Glass

## Abstract

Micro-ribonucleic acids (miRNAs) are key post-transcriptional regulators of the immune system and may play a role in Chronic Obstructive Pulmonary Disease (COPD). In this paper, we constructed subject-specific miRNA-mRNA regulatory networks using bulk and deconvoluted whole blood RNA-sequencing, whole blood miRNA-sequencing, and B-cell receptor-sequencing data from up to 570 miRNAs, 11,859 mRNAs, and 3,190 participants in the COPDGene study.

Analysis of whole blood networks revealed two subnetworks of miRNA-mRNA interactions significantly (FDR<0.05) associated with changes in FEV_1_/FVC. We found that miRNAs (and mRNAs) in the network-identified groups had distinct expression patterns, with miRNAs (and mRNAs) in one group having overall higher expression in COPD (decreasing FEV_1_/FVC) and miRNAs (and mRNAs) in the other group having overall higher expression in controls (increasing FEV_1_/FVC). In addition, miRNAs (and mRNAs) within the same group were positively correlated, while those in different groups were negatively correlated, indicating distinct functional roles for these miRNAs (and mRNAs) as a function of increased COPD severity. Network analysis also identified *PAX5*, a transcription factor master regulator of B-cell development, as the main mRNA network hub. Using ChIP-seq data in lymphoblastoid cells, we identified a PAX5 binding site overlapping with a COPD genome-wide association signal in the promoter region of *ADAM19*. We also found a loss of co-expression between *PAX5* and *ADAM19* in COPD subjects. Furthermore, in B-cell deconvoluted data, *PAX5* was differentially co-expressed with genes associated with B-cell activation and differentiation, revealing a possible mechanism for the regulation of the immune response in COPD. Finally, in B-cell receptor sequencing data, *PAX5* and the identified mRNA subnetworks were negatively associated (FDR<0.05) with immunoglobulin class switching, and positively associated with IgM and IgD counts.

In conclusion, *PAX5* is a known regulator of B-cell identity. B cells are recognized as key players in chronic inflammation and immune dysregulation in COPD. Our work suggests that *PAX5* plays a mediating role both in *ADAM19* regulation and in miRNA regulation of early B cells in COPD.

## Introduction

Chronic Obstructive Pulmonary Disease (COPD), the third leading cause of death worldwide ^1^, is a complex disease whose pathobiology includes airway inflammation ^2^. COPD results from repeated lung insults by toxic exposures, such as cigarette smoking and pollution ^1^, as well as genetic risk factors, many of which have been identified by GWAS (Genome Wide Association Studies) ^3^. Some disease risk is explained by common variant genetics via GWAS; however, the mechanisms of GWAS risk and the molecular drivers of disease pathogenesis independent of genetic risk are unclear. Recent studies have pointed to miRNAs as potential key mediators of the regulatory network associated with the immune response to foreign pathogens ^4^.

Micro-ribonucleic acids (microRNAs, or more simply, miRNAs) are small, non-coding RNA molecules, typically 20-22 nucleotides in length, that play a role in post-transcriptional gene regulation. They bind to complementary sequences on target messenger RNA (mRNA) molecules, usually in the 3′ untranslated region (3′UTR), often resulting in mRNA degradation or the inhibition of translation. miRNA expression is significantly influenced by cigarette smoking ^5,6^, and in macrophages it has been shown that smoking lowers overall miRNA expression ^7^. miRNAs also play a central role in the response to infections and injuries, modulating the immune and physiological processes involved in pathogen removal and maintaining homeostasis ^8^. Specific miRNAs have already been associated with lung diseases, such as COPD, asthma, lung cancer and pulmonary fibrosis^9^. Only a few studies have looked at miRNA regulation in peripheral blood immune cells ^10^. Blood miRNA signatures have been used to identify COPD patients ^11^.

In the lungs of COPD patients, environmental insults lead to the recruitment and activation of local and peripheral B cells, which expand in number and form lymphoid follicles. These activated B cells release cytokines and antibodies, contributing to lung tissue degradation and inflammation, ultimately exacerbating the progression of emphysema and air space enlargement ^12^. B cells at all stages are crucial players in the adaptive immune response to foreign pathogens and have already been associated with COPD severity ^13,14^. In this work we explore the role of miRNA regulation in peripheral blood and specifically B cells ^15,16^. Our goal is to characterize a regulatory mechanism that may modulate the recruitment and activation of peripheral B cells in COPD patients.

The relative redundancy of complementary sequences between miRNAs and their target mRNAs allows a single miRNA to regulate tens to hundreds of genes simultaneously ^17^. Consequently, network analysis can provide a framework for understanding the interaction of multiple miRNAs regulating mRNA levels^18,19^. Networks are composed of nodes, such as miRNA or mRNA molecules, and of edges linking these nodes based on a defined relationship, such as a regulatory interaction – in this case the network can be referred to as a “regulatory network”. Networks provide a natural framework to have a more complete and less biased view of the molecular interactions that make up biological pathways and cellular processes. Therefore, the insights gained from networks can help to expand our understanding of complex diseases, where multiple factors are involved in the pathogenesis.

In this work we investigate miRNA-mRNA regulatory networks in COPD. We leverage two algorithms, LIONESS ^20^ and PUMA ^21^, to infer subject-specific miRNA-mRNA regulatory networks in both whole blood and B-cell deconvoluted data (**Figure 1**). We then analyze these regulatory networks to identify miRNA-mRNA regulatory interactions that are significantly associated with airflow limitation, as measured through FEV_1_/FVC. The identified regulatory interactions point to *PAX5*, a transcription factor and master regulator of B cells, as the most dysregulated mRNA. Finding *PAX5* as a potential key player in the commitment and activation of B cells in the context of COPD paves the way for a deeper understanding of how both *PAX5* and other genes are regulated by miRNAs, their role in blood B cells, and ultimately COPD pathogenesis.

**Figure 1:**
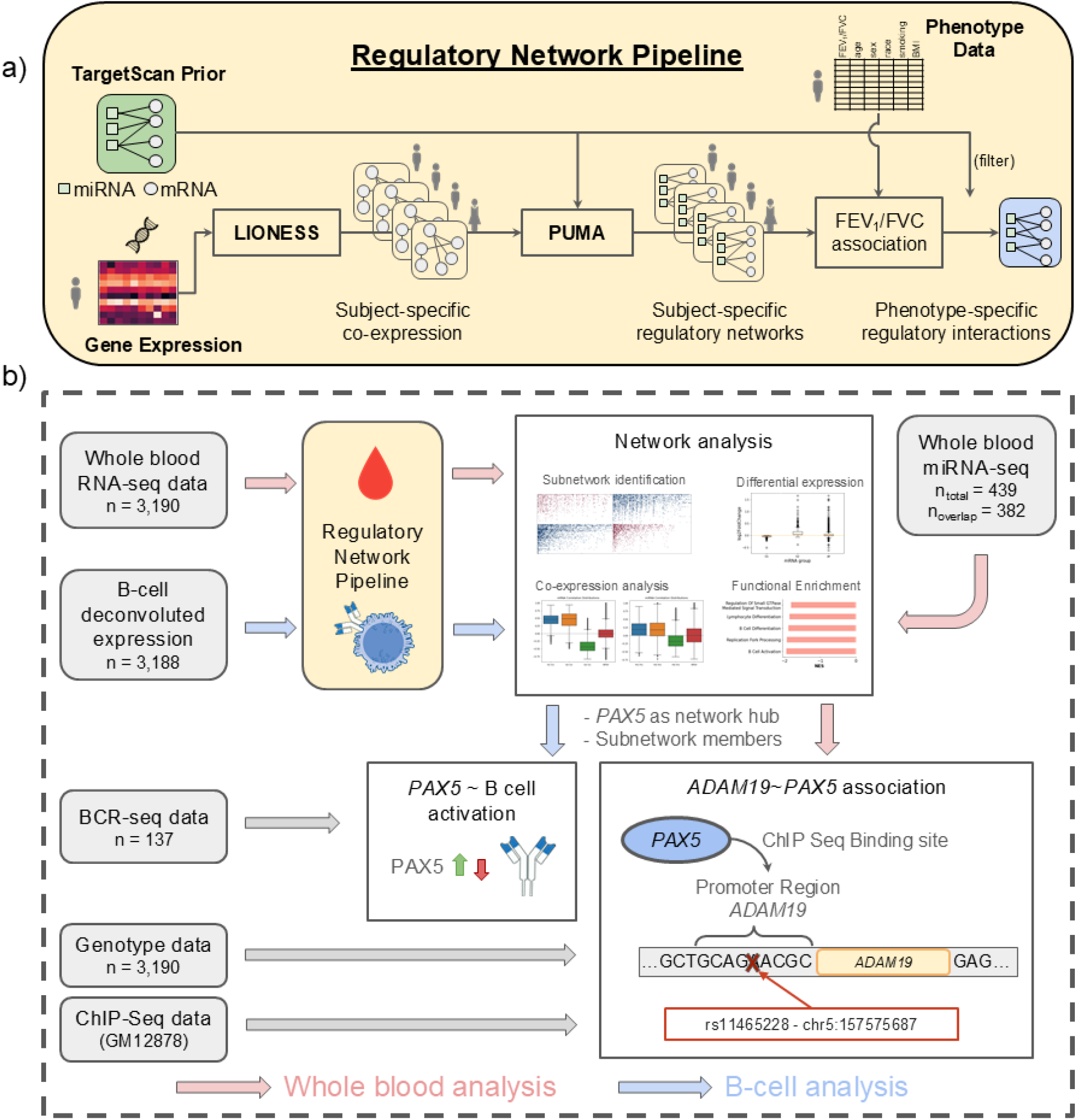
Study design. a) Regulatory Network Inference Pipeline: We computed subject-specific miRNA-mRNA regulatory networks by combining the LIONESS and PUMA algorithms. With LIONESS we inferred subject-specific co-expression from COPDGene RNA-seq data. Then we used PUMA to integrate these data with an initial set of predicted miRNA-mRNA interactions (TargetScan Prior) and obtained subject-specific miRNA-mRNA regulatory networks. Finally, we identified COPD-specific miRNA-mRNA regulatory interactions by linearly associating the PUMA score for each miRNA-mRNA edge in the regulatory networks with FEV_1_/FVC. b) The study is organized around the results from two main analyses performed using whole blood (red arrows) and b-cell (blue arrows) deconvoluted gene expression data. Both analyses start with the inference and analysis of subject-specific miRNA-mRNA networks from gene expression. In both network analyses we found *PAX5* as a key hub and identified two groups of miRNAs and two groups of mRNAs. *PAX5* and these groups served as input for an in-depth analysis of *ADAM19*, which we performed using genotype data, publicly available ChIP-seq data, and B-cell receptor (BCR) data. The miRNA-seq, mRNA-seq, BCR-seq, and genotype data are from subjects in COPDGene study; unless noted otherwise, the number of subjects from each data source are a subset of the 3,190 subjects for which we have whole blood gene expression.

## Materials and Methods

### COPDGene RNA-seq and miRNA-seq expression data

We leveraged whole blood RNA sequencing (RNA-seq) data (n = 3,985) from Phase 2 (5-yr follow-up visit) of the COPDGene study, a longitudinal multicenter observational study of COPD, that was previously collected (clinicaltrial.gov identifier: NCT00608764) and processed as described in Ryu et al 2023 ^22^. We subsequently removed batches with less than 10 subjects as well as PRISm (Preserved Ratio Impaired Spirometry; FEV_1_/FVC >= 0.7 but FEV_1_< 80%) and never smoking subjects, for a final count of 3,190 subjects. We also only kept “protein coding” annotated genes (gencode v37) and those with an associated target mRNA in our TargetScan prior (see below), resulting in a final data set containing 3,190 subjects and 11,859 genes.

We used previously obtained and processed whole blood miRNA sequencing (miRNA-seq) data (n = 538) from Phase 2 of the COPDGene study, where single multiplexed blocking oligonucleotides were used to reduce unwanted hemolysis-related miRNAs hsa-miR-486-5p, hsa-miR-451a, and hsa-miR-92a-3p ^23^. We then followed a similar preprocessing as in Zhuang et. al. ^19^ to remove low quality samples and lowly expressed miRNAs, resulting in a final data set containing 439 subjects and 679 miRNAs.

For additional details on the RNA-seq and miRNA-seq expression data processing, see **Supplemental Materials and Methods**.

### miRNA target prediction data

We leveraged publicly available information on miRNA targeting of mRNAs from TargetScan ^24^ to construct an input motif prior for the PUMA algorithm. We filtered these data to only include miRNAs and mRNAs (genes) expressed in the whole blood miRNA-seq and RNA-seq data, respectively, resulting in a network that included 670 miRNAs, 11,859 mRNAs, and 1,779,028 edges. To avoid high redundancy in our network we then merged miRNAs belonging to the same family into a single node (see **Supplemental Table 1**). This resulted in a network composed of 570 miRNAs, 11,859 mRNAs, and 1,496,256 edges. We weighted the edges in this network by −1 times the TargetScan score, since the PUMA algorithm treats more positive values with higher confidence. We refer to this network as the TargetScan prior. For additional details on processing the miRNA target prediction data, see **Supplemental Materials and Methods**.

### B-Cell Deconvoluted Data

Deconvoluted data were obtained from Ryu et al 2023 ^22^. In this paper, the authors applied CIBERSORTx to deconvolute COPDGene bulk blood RNA-seq data from former and current smokers and impute cell type-specific gene expression for 10 major leukocyte types, including B cells. For our analysis, we selected the deconvoluted “B-cell” gene expression data and filtered it to retain subjects used in our bulk RNA-seq analysis and genes with an associated mRNA target in our TargetScan prior (see above), resulting in B cell gene expression for 3,188 subjects and 2,467 genes.

### B-cell Receptor Sequencing Data

We leveraged adaptive immune receptor repertoire sequencing data for B cell receptors (hereafter, ‘BCR-seq’) that had previously been generated using bulk RNA-seq data from COPDGene^25^. For additional details on BCR-seq data, see **Supplemental Materials and Methods**. We identified 137 subjects with BCR-seq data that also were included in our bulk RNA-seq analysis (see above).

## Statistical Association

We used the python package *statsmodels* (version 0.14.1) to compute the association between BCR features and the expression of genes identified from our network analysis (including *PAX5*, see below) in the B-cell deconvoluted data. We adjusted for age, sex, race, BMI, and smoking status. The full model was:

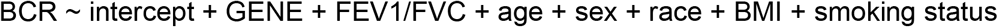

### Regulatory Network Inference

We used a combination of LIONESS ^20^ and PUMA ^21^ to estimate subject-specific miRNA-mRNA regulatory networks (**Figure 1a**). In these networks, nodes are miRNAs and mRNAs and edges represent regulation of the mRNA by the miRNA.

*LIONESS* defines sample-specific networks as *e*^(*q*)^ = *N*(*e*^(*α*)^ *− e*^(*α−q*)^) + *e*^(*α−q*)^, where *e*^(*α*)^ is a network calculated on all samples, *e*^(*α−q*)^ is a network calculated using all but one sample (*q*), *e*^(*q*)^ is the network for sample *q*, and *N* is the total number of samples. We applied LIONESS to obtain sample-specific networks based on Pearson correlation. For the blood RNA-seq data, this resulted in 3,190 subject-specific co-expression networks containing 11,859 genes. For B-cell deconvoluted data, this resulted in 3,188 subject-specific co-expression networks containing 2,467 genes.

*PUMA* estimates miRNA-mRNA regulatory networks by iteratively updating the weights of a prior regulatory network between miRNAs and mRNAs. The algorithm’s objective is to increase the miRNA-mRNA network’s concordance with mRNA (gene) co-expression patterns; miRNA expression information is not considered by the algorithm. *PUMA* models the regulatory network as a bipartite graph, with miRNAs on one side and target mRNAs (genes) on the other. In our analysis, we ran *PUMA* for each subject using two inputs: 1) the TargetScan prior and 2) the individual subject’s LIONESS-inferred co-expression network. As output, we obtained a set of subject-specific miRNA-mRNA regulatory networks (**Figure 1a**). Specifically, for the blood RNA-seq data we obtained 3,190 subject-specific regulatory networks, each containing 570 miRNAs and 11,859 mRNAs. For the B-cell deconvoluted data, before running PUMA we filtered the TargetScan prior to only contain mRNAs expressed in these data. This resulted in 3,188 subject-specific regulatory networks, each containing 570 miRNAs and 2,467 mRNAs. For both the PUMA and LIONESS algorithms, we used the default parameters as implemented in the python package *NetZooPy* ^26^.

### Regulatory Network (PUMA score) analysis

Upon obtaining subject-specific networks, we used the *statsmodels* python package (v 0.14.2) to perform linear regression on the LIONESS-PUMA-estimated miRNA-mRNA edge scores with the following model:

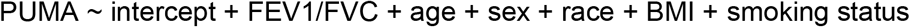

We identified edges whose weight changes significantly (FDR <0.05, Benjamini/Hochberg) as a function of FEV_1_/FVC, adjusting for age, sex, race, BMI, and smoking status. The FEV_1_/FVC ratio is a critical metric for studying COPD as it serves a diagnostic purpose, with COPD typically being defined when FEV1/FVC falls below 0.7. As a continuous variable, it enables the assessment of COPD across a spectrum of disease severities, including mild cases (GOLD stage equal to 1), which are often excluded from studies.

For the networks derived from bulk RNA-seq data, we identified 37,600 significant interactions between 322 miRNAs and 2,460 mRNAs. For the networks derived from B-cell deconvoluted expression data, we obtained 139,913 significant interactions between 376 miRNAs and 2,375 mRNAs. For each of these sets of significant interactions, we subsequently filtered to retain only those that were in the TargetScan prior. Then, we iteratively filtered nodes (miRNAs or mRNAs) until we retained only those nodes with at least *k* of these filtered significant interactions (the k-core of the network). For the networks derived from bulk data, we set *k*=3 (to remove nodes with only 1 and 2 interactions), this identified a total of 7,685 filtered significant interactions between 162 miRNAs and 516 mRNAs; for the networks derived from the B-cell deconvoluted expression data, given the higher number of edges we set *k*=5, this identified a total of 26,199 significant miRNA-mRNA interactions between 265 miRNAs and 776 mRNAs.

We used this information to create a corresponding bipartite network whose edges can take one of three values: −1 if the edge is significant and increases in weight with *decreasing* FEV_1_/FVC (“COPD-specific”); 1 if the edge is significant and increases in weight with *increasing* FEV_1_/FVC (“control-specific”); 0 otherwise. We then clustered this bi-partite network using agglomerative clustering on the Hamming distance between pairs of nodes (mRNAs and miRNAs separately) using the AgglomerativeClustering (affinity = ‘precomputed’, linkage=‘average’) function from the *sklearn* python package v1.5.0.

### PAX5 co-expression analysis

The main mRNA network hub identified from the above analysis is *PAX5*. Therefore, for both for bulk and B-cell deconvoluted data, we next computed the partial correlation of *PAX5* with mRNAs and miRNAs separately in COPD cases and control subjects. We defined COPD subjects as those with FEV1<80% and FEV1/FVC<0.7 and control subjects as those with FEV1>=80% and FEV1/FVC>=0.7, we did not consider other subjects (i.e. those with GOLD stage equal to 1) in this analysis in order to maximize the separation between COPD cases and controls. We used partial correlation to compute the co-expression while accounting for possible confounding factors. For the bulk data we adjusted for age, sex, race, BMI, smoking status, and cell composition (white blood cell counts, lymphocyte percentage). In the B-cell deconvoluted data, since the cell composition data have been used to infer cell-specific gene expression, we only adjusted for age, sex, race, BMI, and smoking status.

### Differential expression analysis

For both the miRNA-seq and RNA-seq data, we used the python package *pydeseq2* (version 0.4.9) to compute the differential expression of each miRNA (and mRNA) using FEV_1_/FVC as the target variable and batch, age, sex, race, BMI, and smoking status as confounding factors. For whole blood expression data, in order not to be biased by cell composition, we also performed analyses that added white blood cell counts and lymphocyte percentage as confounding factors.

### LIONESS differential co-expression and GSEA pre-Rank enrichment analysis

Differential co-expression between gene pairs is usually computed with respect to two or more discrete conditions ^27,28^, such as between COPD cases and controls. Here we take advantage of the subject-specific LIONESS scores to model how co-expression changes as a function of a continuous outcome, in our case FEV_1_/FVC. Specifically, for each gene, we computed the linear association between the subject-specific co-expression values of that gene with *PAX5* (as computed by LIONESS) and FEV_1_/FVC. We then used these ‘differential co-expression values’ to pre-Rank genes and perform Gene Set Enrichment Analysis (GSEA). GSEA was performed using the python package *gseapy* (version 1.1.3). We used the gene sets from the Biological Process of Gene Ontology (GO) ^29^, namely “*GO_Biological_Process_2023*”. We selected sets with min_size=10 and max_size=50. We set the permutation_num=10000 and seed=6.

### PAX5-ADAM19 association

#### Chip-seq data

We downloaded Chip-seq data for hg38 non-redundant peaks as defined by ReMap2022 ^30^ (https://remap.univ-amu.fr/download_page; accessed August 10, 2021). From this file we extracted all peaks for *PAX5* in GM12878 (a lymphoblastoid cell line). We then used the bedtools intersect function (bedtools v2.29.2) to identify *PAX5* peaks that overlap with gene promoter regions, defined as [−250,+50] around gencode v37 annotated transcriptional start sites, for a total of 15,901 unique *PAX5* binding sites located in 9794 gene promoter regions; 5847 of these genes correspond to mRNAs in our whole blood network model.

#### COPD GWAS data

COPD GWAS data were collected from two previous studies, Shrine et. al.^31^ and Sakornsakolpat et. al.^32^. We selected GWAS SNPs associated with COPD or FEV_1_/FVC for a total of 556 SNPs. We identified which of these SNPs fall within *PAX5* binding sites located in gene promoter regions, as defined above. **Supplemental Table 2** shows the results of this intersection and reports the overlapping SNPs’ rsIDs and genomic (GRCh38) locations. We used statistical fine mapping to identify credible sets and the most likely causal variants for these SNPs. For COPD associated SNPs, we used the approximate Bayes factor calculation as detailed in Sakornsakolpat et. al.^32^, and for FEV_1_/FVC SNPs from Shrine et. al., we used European ancestry summary statistics and the SuSiE (Sum of Single Effects) model ^33^ as implemented in the R package *susieR* (version 0.12.35). We conducted fine mapping with a window of 2×10^5^ base pairs centered at the lead variant. We removed variants that were duplicated or that had a low sample size, defined as the lower quartile of all sample sizes in the data.

#### Statistical Association

We used the python package *statsmodels* (version 0.14.1) to compute the association between *ADAM19* and *PAX5* expression in the bulk COPDGene expression data, treating *ADAM19* expression as the outcome, and including an interaction term between FEV_1_/FVC and *PAX5* expression, as well as an interaction term between rs11465228 (GWAS variant in *ADAM19* promoter) and *PAX5* expression. We also adjusted for white blood cell counts, lymphocyte percentage, age, sex, race, BMI, smoking status, and the first 10 principal components of genetic ancestry. The full model was:

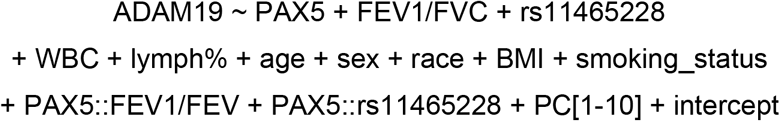

## Results

### miRNA-mRNA regulatory interactions associated with FEV_1_/FVC

We leveraged a subpopulation of the COPDGene Study (**Table 1**) with RNA-seq data collected during the 5-year follow-up visit (Phase 2). These data include gene expression profiles for 3,190 subjects and 11,859 genes (mRNAs). We inferred miRNA-mRNA regulatory interactions significantly (FDR < 0.05) associated with FEV_1_/FVC and filtered to retain a core network of 7,685 significant interactions between 162 miRNAs and 516 mRNAs. The degree (number of filtered significant interactions identified) for the top 50 mRNA and miRNA is reported in **Supplemental Tables 3-4**.

**Table 1:** Clinical Characteristics of the Study Population. P-values were computed with ANOVA test for continuous variables and Chi-squared test for categorical variables.

|  |  | **GOLD 0** | **GOLD 1** | **GOLD 2** | **GOLD 3** | **GOLD 4** | **P-Value** |
| --- | --- | --- | --- | --- | --- | --- | --- |
| **n** |  | 1607 | 366 | 708 | 361 | 148 |  |
| **Age, mean (SD)** |  | 63.4 (8.3) | 68.8 (8.6) | 67.6 (8.6) | 69.1 (8.1) | 67.6 (7.7) | <0.001 |
| **BMI, mean (SD)** |  | 29.2 (6.0) | 26.6 (4.6) | 28.8 (6.3) | 27.8 (6.6) | 26.0 (6.1) | <0.001 |
| **FEV1 FVC post, mean (SD)** |  | 0.78 (0.05) | 0.65 (0.04) | 0.59 (0.08) | 0.44 (0.09) | 0.34 (0.08) | <0.001 |
| **FEV1 FVC pre, mean (SD)** |  | 0.8 (0.1) | 0.6 (0.1) | 0.6 (0.1) | 0.4 (0.1) | 0.3 (0.1) | <0.001 |
| **Duration Smoking, mean (SD)** |  | 34.9 (11.2) | 39.7 (11.7) | 40.9 (10.5) | 42.7 (9.8) | 41.0 (8.9) | <0.001 |
| **gender, n (%)** | **Female** | 851 (53.0) | 154 (42.1) | 320 (45.2) | 151 (41.8) | 62 (41.9) | <0.001 |
|  | **Male** | 756 (47.0) | 212 (57.9) | 388 (54.8) | 210 (58.2) | 86 (58.1) |  |
| **race, n (%)** | **Black** | 507 (31.5) | 74 (20.2) | 158 (22.3) | 71 (19.7) | 26 (17.6) | <0.001 |
|  | **White** | 1100 (68.5) | 292 (79.8) | 550 (77.7) | 290 (80.3) | 122 (82.4) |  |
| **smoking status, n (%)** | **Current** | 614 (38.2) | 139 (38.0) | 281 (39.7) | 97 (26.9) | 24 (16.2) | <0.001 |
|  | **Former** | 993 (61.8) | 227 (62.0) | 427 (60.3) | 264 (73.1) | 124 (83.8) |  |

Clustering of these interactions (based on pair-wise Hamming distance) indicated that the network formed by the obtained set of significant interactions is highly modular (**Figure 2a**), and made up of two subnetworks that are composed of two distinct groups of miRNAs, m1 (n=82) and m2 (n=80), and two distinct groups of mRNAs, G1 (n=254) and G2 (n=262). Significant regulatory interactions between miRNA and mRNA groups are predominantly either positively (blue dots) or negatively (red dots) associated with FEV_1_/FVC. Robustness analysis of these clustering results to network preprocessing and input randomization are found in **Supplemental Figure 1** and **Supplemental Materials and Methods**.

**Figure 2:**
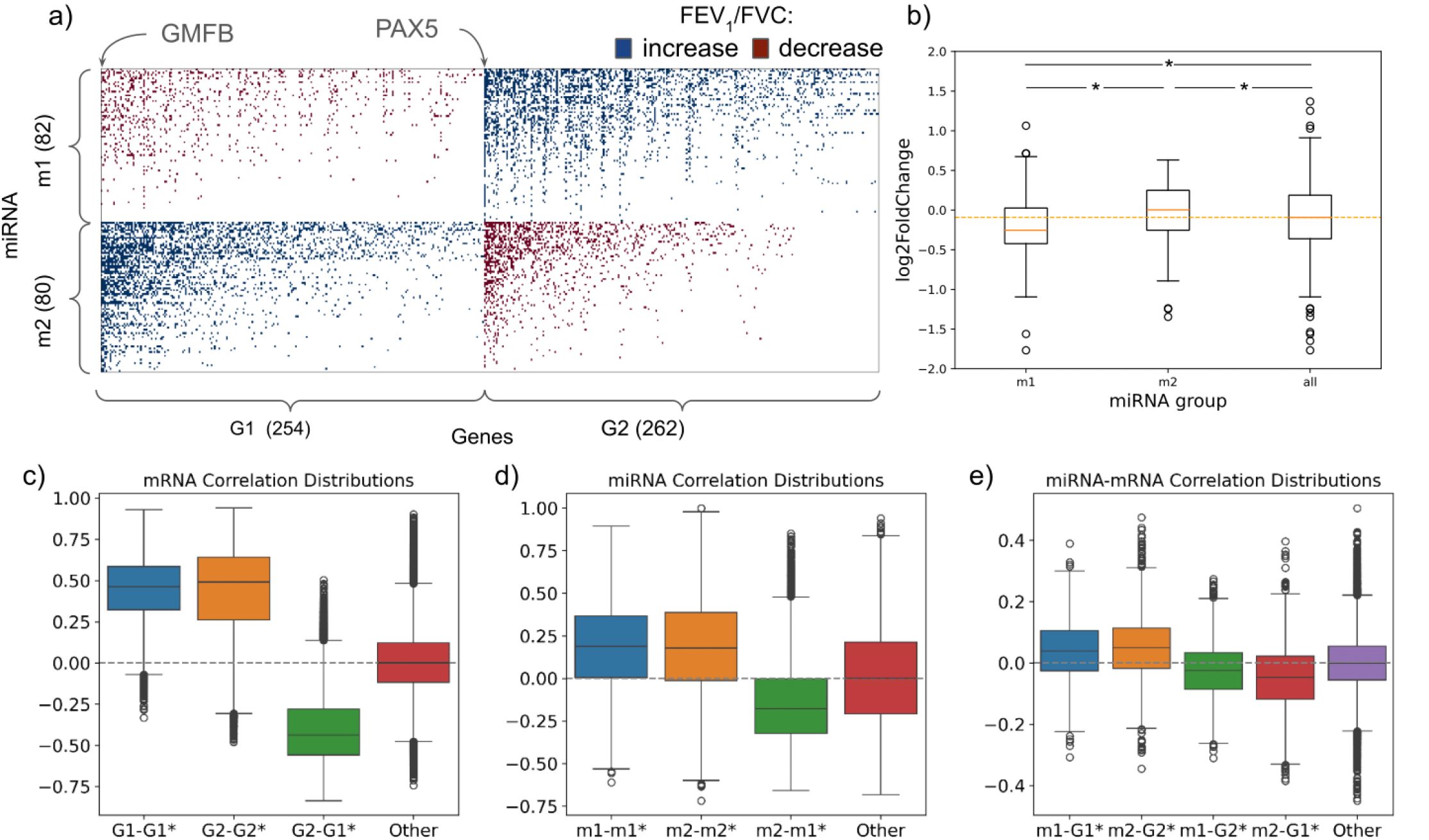
Network analysis using bulk whole blood expression data. **(a)** The clustered adjacency matrix of filtered miRNA-mRNA interactions significantly associated with FEV_1_/FVC. A dot in the matrix represents a significant edge between the miRNA (row) and mRNA (column). The color of the dot represents the sign of the association. Rows and columns have been sorted according to their degree and their subnetwork group (m1, m2, G1, G2). **(b)** Boxplots showing the distribution in log2 fold change values for miRNAs in m1 and m2 when computing the differential expression with respect to changes in FEV_1_/FVC. An asterisk (*) indicates p<0.05 by two-sided t-test. **(c-e)** Boxplots showing the distribution in for **(c)** mRNA, **(d)** miRNA, and **(e)** miRNA-mRNA pairwise correlations within and between subnetwork groups. For miRNA-mRNA pairwise correlations, we only show those that had a significant filtered interaction based on our network analysis. The “Other” group is formed by all other mRNA, miRNA, and miRNA-mRNA pairs. An asterisk (*) next to the label indicates p<0.05 by two-sided t-test comparing the distribution of pairwise correlations values in the that group versus “other”.

### mRNA co-expression and differential expression analysis

To evaluate the expression profiles for the mRNAs in G1 and G2, we determined the differential expression of these mRNAs as a function of FEV_1_/FVC and observed significant opposite trends for mRNAs in G1 and G2 (p<0.001). In particular, as illustrated by their negative log2 fold change with respect to increasing FEV_1_/FVC (**Supplemental Figure 2**), mRNAs in G1 are significantly (p<10^−20^, one sided t-test) more expressed in the context of more severe COPD, while mRNAs in G2 tend to be less expressed (positive log2 fold change, p<10^−11^). Next, we calculated the correlation of these mRNAs. We observe that mRNAs that are assigned to the same group are highly correlated, while mRNAs assigned to different groups tend to be anti-correlated (**Figure 2c**).

### miRNA co-expression and differential expression analysis

Next, we leveraged miRNA-seq data collected on 538 subjects in COPDGene and performed an analogous analysis. We do not see strong differential expression of individual miRNAs as a function of FEV_1_/FVC, with only two miRNAs (mir-4488 and mir-6087) significantly differentially expressed (FDR <0.05). However, the log2 fold change of miRNAs is significantly different between miRNAs in m1 and m2 (p<0.005; **Figure 2b**). Furthermore, we once again observe that miRNAs assigned to the same group tend to be highly correlated, while miRNAs assigned to different groups tend to be anti-correlated (**Figure 2d**). Since the expression levels of miRNAs were not used to build the network, these results indicate that our network analysis is correctly predicting biological signals not contained in the input data.

### miRNA-mRNA co-expression analysis

Finally, we leveraged data from an overlapping set of 382 subjects for which we have both miRNA-seq and RNA-seq data and calculated the co-expression between miRNA-mRNA pairs with an identified filtered significant interaction. Interactions that extend from miRNAs in m1 to mRNAs in G1 (m1-G1) as well as from miRNAs in m2 to mRNAs in G2 (m2-G2) tend to be more positively correlated, while interactions from miRNAs in m1 to mRNAs in G2 (m1-G2) as well as from miRNAs in m2 to mRNAs in G1 (m2-G1) tend to be more negatively correlated (**Figure 2e**). Notably, regulatory interactions between m1-G2 and m2-G1 were associated with increasing FEV_1_/FVC (non-COPD), suggesting a possible loss of regulation between these miRNAs and mRNAs in the COPD phenotype. Given the primary role of miRNAs in suppressing mRNA expression, our results suggest these regulatory interactions may be functional.

### PAX5 identified as a hub among FEV_1_/FVC associated interactions

The two most targeted mRNAs in G1 and G2 are *GMFB* and *PAX5*, respectively, both of which are highly involved in B-cell processes ^34–36^. Given this result, we ran a sensitivity analysis in which we repeated our network analysis while controlling for cell composition; we still identified *PAX5* (but not *GMFB*) as a key hub in the network (**Supplemental Figure 3**).

#### PAX5-mRNA co-expression analysis

To interpret this result, we determined the co-expression of *PAX5* with respect to the other mRNAs in G1 and G2; we performed this analysis separately for COPD cases and control subjects and used partial correlation analysis to account for cell composition and potential confounding factors (see **Methods**). In COPD subjects, we observe lower absolute *PAX5* co-expression with mRNAs in G1 (less negatively co-expressed) and G2 (less positively co-expressed) (**Figure 3a**). This may indicate a potential loss of PAX5 regulation of these mRNAs in the context of COPD.

**Figure 3:**
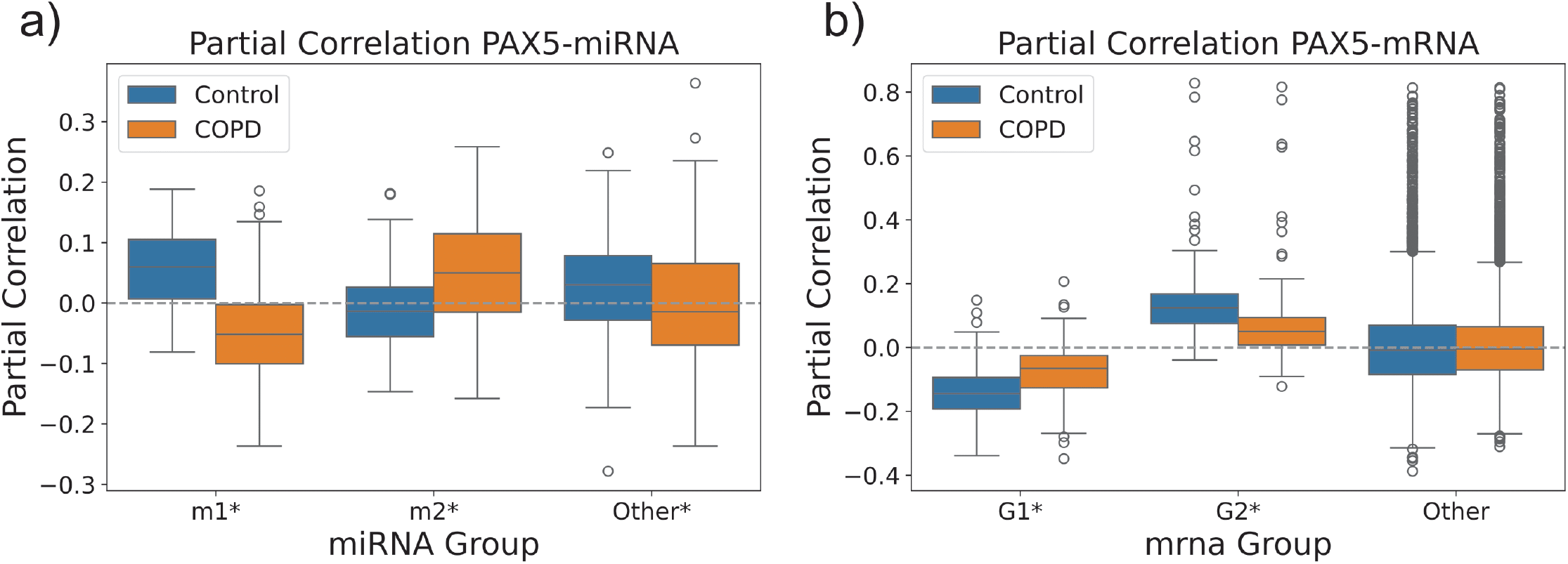
Partial Correlation of network-identified mRNAs and miRNAs with PAX5. **(a)** The partial correlation of *PAX5* expression with other mRNAs in whole blood RNA-seq data, calculated separately for COPD cases and controls. The distribution of these correlation values is shown separately for mRNAs in G1, mRNAs in G2, and other mRNAs. **(b)** The distribution of the partial correlation of *PAX5* expression with miRNAs, calculated separately for COPD cases and controls, and shown separately for miRNAs in m1, miRNAs in m2, and other miRNAs. Note the “Other” group in each plot represents a random set of mRNAs or miRNAs, respectively, which has been matched in size to the largest of two groups (G2 and m1). An asterisk (*) next to the label indicates p<0.05 by two-sided t-test comparing the distribution of partial correlation values in COPD versus control.

#### PAX5-miRNA co-expression analysis

Next, we evaluated the co-expression of *PAX5* with respect to the miRNAs in m1 and m2, using the same approach as with the mRNAs. We find strong differential co-expression patterns for *PAX5* with m1 miRNAs (**Figure 3b**). PAX5 is generally positively correlated with m1 miRNAs across control subjects. However, the opposite is true in COPD, with PAX5 negatively correlated with m1 miRNAs in COPD subjects. This suggests a possible change in miRNA regulation of *PAX5* in COPD compared to control subjects. PAX5 is also generally positively correlated with m2 miRNAs in COPD.

#### PAX5 ChIP-seq analysis

*PAX5* is a well-studied transcription factor master regulator of B-cell development. We leveraged publicly available Chip-seq data in a lymphoblastoid cell line (GM12878) ^30^ to investigate the regulatory targets of *PAX5* in the context of our network analysis, specifically focusing on *PAX5* binding sites within the promoter regions of genes. We found that genes that correspond to mRNAs in G2 (but not G1) are enriched (p=1.9×10^−4^, Fisher’s exact test) for *PAX5* regulatory targeting; 158 out of the 262 mRNAs in G2 have a validated Chip-seq *PAX5* binding site in their associated gene promoter region.

Next, we evaluated whether *PAX5* binding sites directly overlapped with any known COPD GWAS variants. We found a direct overlap with six COPD GWAS SNPs ^30,31^ (**Supplemental Table 2**). For each of these COPD GWAS SNPs, we identified their corresponding *PAX5*-regulated gene (based on the promoter in which the overlap was located). For these genes, we determined their co-expression with *PAX5* (see **Methods**) separately for COPD cases and controls (**Table 2**). For rs11465228, its corresponding PAX5-regulated gene, *ADAM19*, was observed to be positively correlated with *PAX5* in controls, but this correlation significantly decreased in COPD (p=0.005), suggesting a possible loss of regulation of *ADAM19* by *PAX5* in the context of COPD. It is worth noting that there are multiple lines of evidence pointing to *ADAM19* being the functional gene associated with rs11465228, including eQTL, mQTL and significant chromatin interaction analysis ^32^.

**Table 2:**
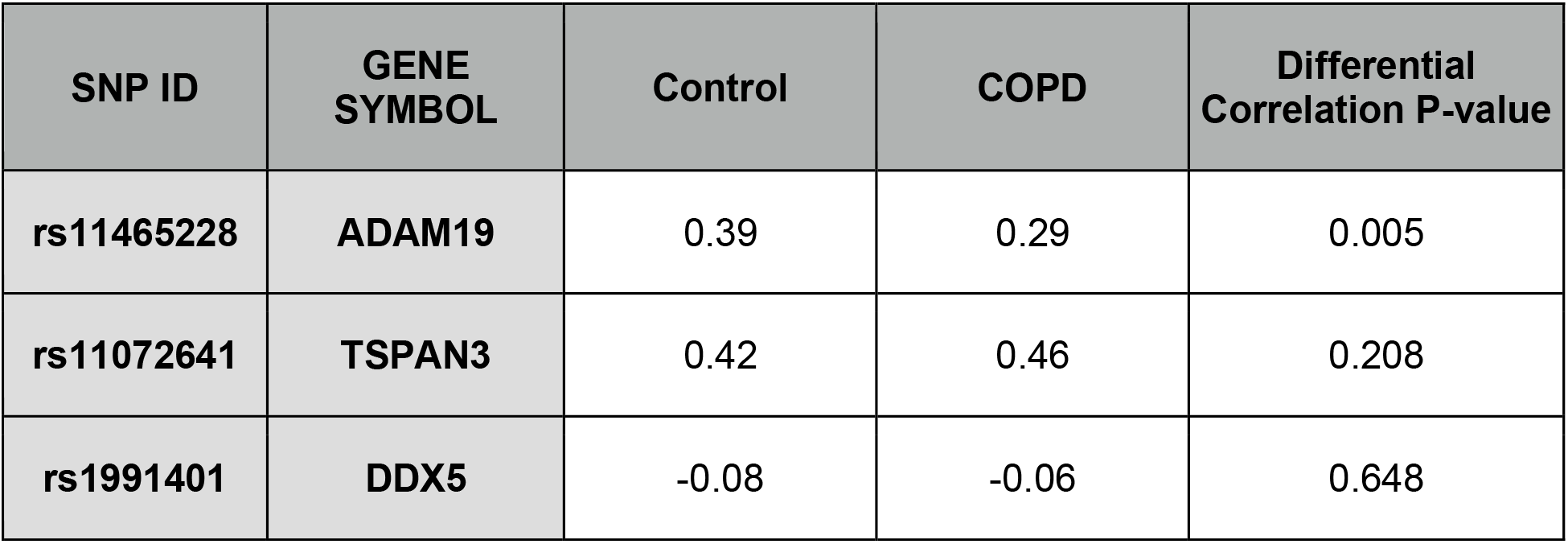
PAX5-gene co-expression in COPD cases and controls. The genes identified by overlapping COPD GWAS SNPs with PAX5 binding sites in gene promoters. The co-expression between *PAX5* and each of the genes was obtained by computing the partial correlation on the whole blood RNA-seq from COPDGene and adjusting for white blood cell counts, lymphocyte percentage, age, sex, race, BMI, and smoking status. P-values were computed based on the z-score of the Fisher transformation.

| SNP ID | GENE SYMBOL | Control | COPD | Differential Correlation P-value |
| --- | --- | --- | --- | --- |
| rs11465228 | ADAM19 | 0.39 | 0.29 | 0.005 |
| rs11072641 | TSPAN3 | 0.42 | 0.46 | 0.208 |
| rs1991401 | DDX5 | -0.08 | -0.06 | 0.648 |

Furthermore, rs11465228 is the most likely causal variant at the locus for both COPD and lung function, with a posterior probability of being the causal variant of > 80% for FEV_1_/FVC ratio (lung function).

### Association of ADAM19 with PAX5 and miRNA expression levels

To further understand the relationship between *PAX5* and *ADAM19*, we ran a linear model quantifying the statistical association between their expression across subjects while accounting for other relevant variables, including FEV_1_/FVC, the rs11465228 genotype, white blood cell count, lymphocyte percentage, age, sex, race, BMI, and smoking status; the model also included two interaction terms: (1) between *PAX5* expression and FEV_1_/FVC, and (2) between PAX5 expression and the rs11465228 genotype (**Table 3;** see **Methods** for more details). *PAX5* expression is highly positively associated with *ADAM19* expression (p=7.51×10^−89^). Interestingly, the interaction between *PAX5* and FEV_1_/FVC is also significantly positively associated with *ADAM19* expression (p=6.16×10^−12^), suggesting that the association between *PAX5* and *ADAM19* is negatively disrupted by increased airflow obstruction. The most significantly associated variable was lymphocyte percentage (p=7.51×10^−163^). As expected, the rs11465228 genotype is also significantly positively associated with *ADAM19* expression (p=9.18×10^−8^).

**Table 3:** *PAX5*-*ADAM19* linear association results. The table shows the statistics of the most significantly associated variables from the linear association model between *PAX5* and *ADAM19* expression. The first column of the table indicates the variable of the linear model, “:” means interaction.

| Variable | Coefficient (std) | P-value |
| --- | --- | --- |
| Lymphocyte (%) | -0.53 (0.02) | $8.35 \times 10^{-163}$ |
| PAX5 | 0.37 (0.02) | $1.42 \times 10^{-89}$ |
| PAX5:FEV1/FVC | 0.10 (0.01) | $6.52 \times 10^{-12}$ |
| rs11465228 | 0.21 (0.04) | $9.27 \times 10^{-8}$ |

*PAX5* was identified as a hub highly targeted by many miRNAs. To determine if these miRNAs are also associated with *ADAM19* expression, we identified COPDGene subjects with both miRNA-seq and mRNA-seq data. We then used these data to run a series of linear models evaluating the association of *ADAM19* expression with the expression of each of the miRNAs in the m1 and m2 groups; these models included the same set of variables and interaction terms as in the previous analysis. We observe that miRNAs in m1 tend to be negatively associated with *ADAM19* expression while miRNAs in m2 are positively associated (**Figure 4**).

**Figure 4:**
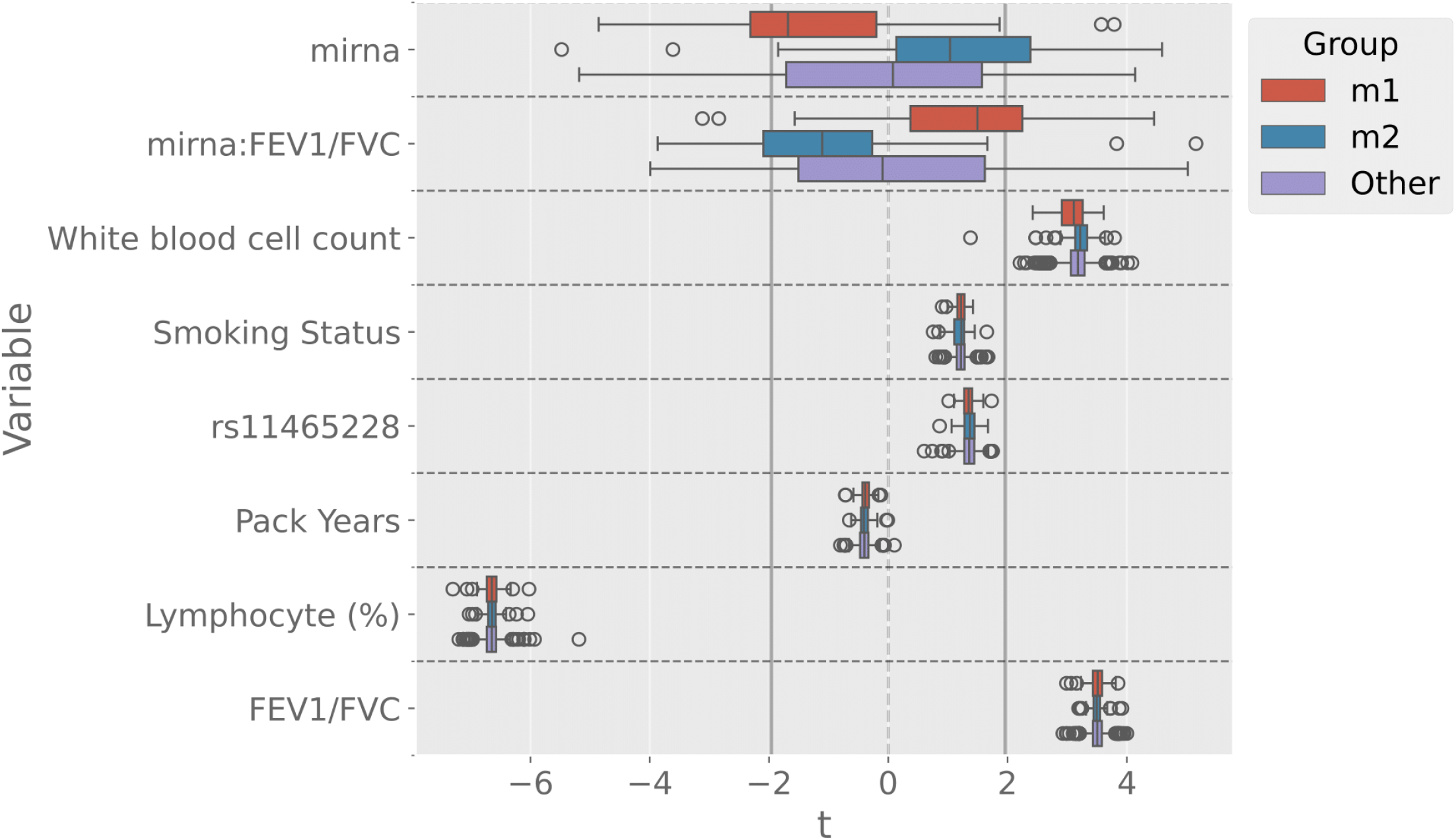
miRNA-*ADAM19* linear association results. The boxplot shows the distribution of *t-statistic* scores of the linear model coefficients associating each miRNA with *ADAM19*. The sign of the *t-statistic* represents the direction of the association, and the absolute value is proportionate to its significance. The vertical black lines represent the negative and positive significance threshold (FDR < 0.05) on the t-statistic.

The interaction term between the miRNAs’ expression levels and FEV_1_/FVC also strongly differs based on miRNA group membership and is in the opposite direction as the miRNA-*ADAM19* expression association, with a positive association for miRNAs in m1 and negative association for miRNAs in m2. This suggests the association between the miRNAs and *ADAM19* is reduced in the context of COPD.

### B-cell specific miRNA-mRNA regulatory network

*PAX5* is a master regulator for B-cell differentiation and specialization ^34,35^. Therefore, we next leveraged B-cell deconvoluted gene expression data from the COPDGene study, obtained from Ryu et al. ^22^. Using the same approach as with the bulk data, we built subject-specific miRNA-mRNA regulatory networks based on this B-cell deconvoluted expression data and identified edges significantly associated with FEV_1_/FVC. We identified 26,199 significant miRNA-mRNA interactions between 265 miRNAs and 776 mRNAs. Filtering and clustering these significant interactions, using the same approach as we applied to the bulk networks, once again revealed a modular structure, with two sets of mRNAs, bG1 (n=109) and bG2 (n=156), and two sets of miRNAs, bm1 (192) and bm2 (584). There is some consistency between the miRNA groups with those identified in the bulk analysis; bm1 and m1 share 54 miRNAs, and bm2 and m2 share 54 miRNAs. In contrast, the mRNA groups only share a handful of genes. However, we once again identify *PAX5* as a hub in bG2. **Figure 5a**, shows the top 20 miRNAs and mRNAs in these groups based on their total number of filtered significant regulatory interactions.

**Figure 5:**
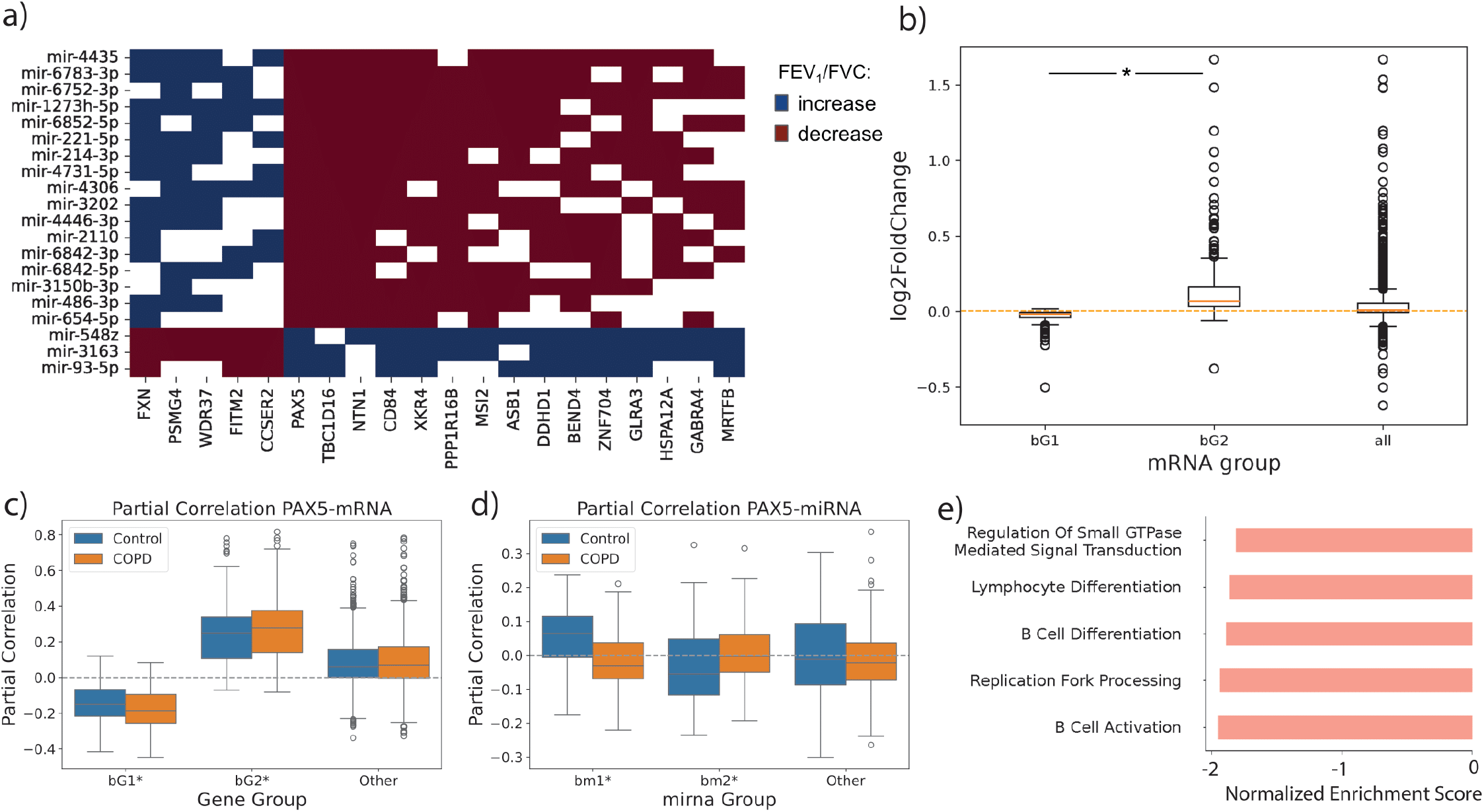
Network analysis using B cell deconvoluted expression data. **(a)** The clustered adjacency matrix of filtered miRNA-mRNA interactions significantly associated with FEV_1_/FVC. Rows and columns have been sorted according to their degree and their subnetwork grouping. Only the top 20 miRNAs and mRNAs based on degree are shown. **(b)** Boxplots showing the distribution in log2 fold change values for mRNAs in bG1 and bG2 when computing the differential expression with respect to changes in FEV_1_/FVC in the B-cell deconvoluted expression data. An asterisk (*) indicates p<0.05 by two-sided t-test. **(c-d)** Boxplots showing the distribution in the partial correlation of PAX5 in the B-cell deconvoluted expression data with the expression of **(c)** mRNAs in bG1 and bG2 (B-cell deconvoluted data) and **(d)** miRNAs in bm1 and bm2 (bulk data); for the miRNAs we only show those that were predicted to target PAX5 based on the TargetScan prior. The “other” group represents the set of mRNAs and miRNAs which are not in either of the other two groups. An asterisk (*) next to the label indicates p<0.05 by two-sided t-test comparing the distribution of partial correlation values in COPD versus control. **(e)** GSEA pre-rank enrichment analysis on mRNAs differentially co-expressed with *PAX5* as a function of FEV_1_/FVC. A negative enrichment score indicates genes in that pathway tend to have increased co-expression with PAX5 with decreased lung function.

We next determined the differential expression of mRNAs with respect to FEV_1_/FVC in the B-cell deconvoluted expression data. As in the bulk data, we observe that the two mRNA groups identified by our network analysis have opposite overall association with FEV_1_/FVC (**Figure 5b**), suggesting a switching behavior with respect to miRNA regulation and the FEV_1_/FVC phenotype. As noted above, *PAX5* is once again one of the most highly targeted mRNAs in our network. Therefore, we evaluated the co-expression of *PAX5* with other mRNAs in the B-cell deconvoluted expression data; as before, we performed this analysis separately for COPD cases and control subjects and used partial correlation in order to account for potential confounding factors (see **Methods**). We find that mRNAs in bG1 tend to be negatively correlated with *PAX5* expression while mRNAs in bG2 tend to be positively correlated; in contrast to our results in the bulk data, the absolute co-expression slightly increases in COPD subjects (**Figure 5c**).

We also determined the partial correlation between *PAX5* expression in the B-cell deconvoluted data with miRNA expression in the bulk data (deconvoluted miRNA expression data was not available). We observe that the correlation of *PAX5* expression with miRNA expression differs between COPD cases and controls for miRNAs in both bm1 and bm2 (**Figure 5d**). In particular, similar to our results in the bulk data, for miRNAs in bm1 we observe a switch from positive to negative correlation with *PAX5* in control versus COPD subjects, respectively. This suggests a change in miRNA regulation of *PAX5* with respect to the COPD phenotype.

Next, to find *PAX5* regulated processes affected by the COPD phenotype, we performed pre-ranked gene set enrichment analysis based on the differential co-expression of mRNAs with *PAX5* with respect to FEV_1_/FVC (see **Methods**). In this analysis, we found a strong enrichment for B cell activation and differentiation, cell-cell junction (necessary for antigen presenting functions) and replication fork processing (highly related to B-cell activation) in the context of decreasing FEV_1_/FEV (**Figure 5e**). These results confirm the regulatory role of *PAX5* in B-cell specific functions and point to its potential role in COPD.

### *PAX5* and mRNAs associated with B-cell activation

Finally, to establish the association between *PAX5* and the mRNAs identified through our network analysis in the context of B-cell specific immune functions, we leveraged a recently generated BCR-seq dataset (see **Methods**) from the COPDGene Study ^25^. 137 subjects had both RNA-seq and BCR-seq data.

To understand whether the mRNAs identified from our network analysis are related to B-cell differentiation and activation, we tested for linear associations between features measured in the BCR-seq data and the expression of mRNAs belonging to the bG1 and bG2 groups (**Figure 6**). Interestingly, many mRNAs in bG2 (including *PAX5*, which is a member of bG2) are significantly positively associated with IGHM and IGHD proportional counts and significantly negatively associated with IGHA and IGHG proportional counts. Many bG2 mRNAs also are significantly negatively associated with class switching. In contrast, bG1 mRNAs have few significant associations with BCR-seq features.

**Figure 6:**
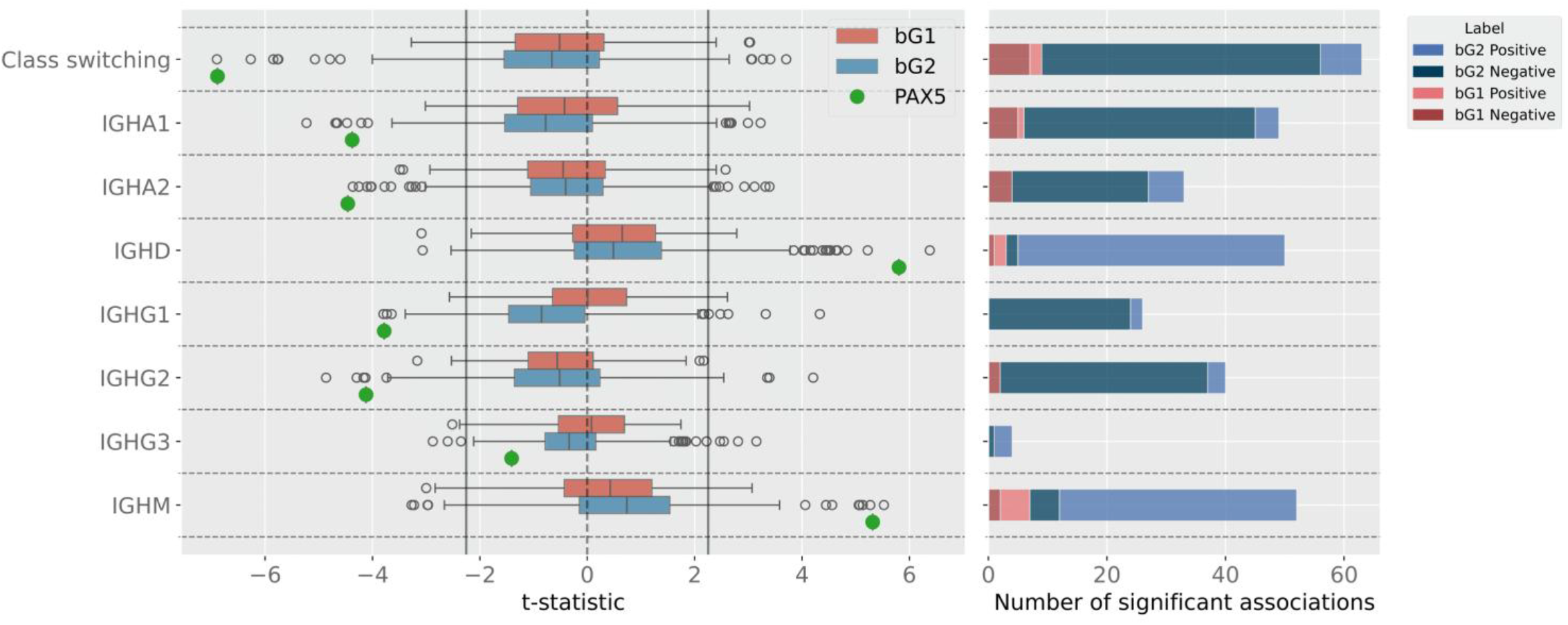
Association of network-identified mRNAs with BCR-seq features. The left plot represents the t-statistic of the linear model association between BCR-seq features and mRNAs in bG1 *(blue)*, bG2 *(red)*, and *PAX5 (green)*, which is a member of bG2. The sign of the *t-statistic* represents the direction of the association, and the absolute value is proportionate to its significance. The vertical black lines represent the negative and positive significance threshold (FDR < 0.05) on the t-statistic. The right plot shows the number of significant associations (FDR < 0.05) based on the mRNA group and whether the associations are positive or negative.

## Discussion

Given the potential role of miRNAs as regulators of the adaptive immune response in COPD, in this paper we studied subject-specific miRNA-mRNA regulatory networks generated from whole blood and deconvoluted B-cell RNA-seq data from a multicenter cohort of smoking subjects with and without COPD. Using these networks, we identified miRNA-mRNA interactions statistically associated with airflow obstruction (FEV_1_/FVC) and found that the subnetwork formed by these interactions had a highly modular structure, with mRNAs and miRNAs falling into distinct groups. Analyzing this subnetwork also identified *PAX5*, a transcription factor and key regulator of innate immunity and B-cell activation, as an mRNA network hub. In investigating the genes regulated by this transcription factor, we identified a direct regulatory interaction between *PAX5* and *ADAM19*, a COPD GWAS gene. Subsequent gene expression analysis indicated that this regulatory interaction is impacted in the context of COPD. Finally, we confirmed *PAX5*’s regulatory role in B-cells by leveraging B-cell receptor data from this same cohort, identifying associations between our network-identified groups and pro-B and pre-B cell activation.

Our network analysis in whole blood identified two groups of miRNAs and two groups of mRNAs. Interestingly, although none of the miRNAs in the network-identified groups are individually significantly differentially expressed with respect to airflow obstruction, collectively miRNAs within groups were shifted towards either increased or decreased expression as a function of FEV_1_/FVC. This result suggests that miRNA regulation occurs at the group level rather than at the individual level. We also found that mRNAs in the network-identified groups had distinct expression patterns, with mRNAs in one group having overall higher expression in COPD (decreasing FEV_1_/FVC) and mRNAs in the other group having overall higher expression in controls (increasing FEV_1_/FVC). In addition, miRNAs and mRNAs were positively co-expressed within groups and negatively co-expressed between groups. Importantly, miRNA expression information was not used to construct the miRNA-mRNA networks. Thus, these findings demonstrate that our network analysis identified biological signals not contained in the input data.

Network analysis also indicated high miRNA targeting of *PAX5*, a transcription factor master regulator of B-cell identity. *PAX5* is necessary in the early stages of B-cell activation, orchestrating the transcriptional programs required for B-cell identity and function. It does this by promoting the expression of genes necessary for B-cell receptor signaling, antigen processing, and presentation, while simultaneously repressing genes associated with alternative lineages to maintain B-cell lineage commitment ^35^. When comparing between COPD cases versus controls, we found that *PAX5* displayed differential co-expression patterns with genes and miRNAs in the network-identified groups. ChIP-seq in lymphoblastoid cells (GM12878) also indicated *PAX5* binding in gene promoters that directly overlapped with the location of six COPD GWAS SNPs, including rs11465228, which is the most likely causal variant associated with *ADAM19*. We found that the expression levels of miRNAs identified by the network, as well as of *PAX5*, were significantly (FDR <0.01) associated with *ADAM19* expression. We also found that the interaction term with FEV_1_/FVC has the opposite direction, implying that the association to ADAM19 decreases in the context of COPD. Overall, these findings suggest that in B cells, the disruptive effect of COPD on *ADAM19* may be mediated by *PAX5* ^37,38^.

*ADAM19* is a human metalloprotease disintegrin associated with EMT (epithelial mesenchymal transition) and immune infiltration ^39^ that is expressed in multiple types of immune cells, including naive, memory, and plasma B cells ^40,41^. In tumors, *ADAM19* has been found to regulate immunotherapy response by releasing signals from the cell surface to improve communication between tumor and B cells ^42^. In the context of COPD, previous studies have demonstrated a functional role for *ADAM19* in the context of alveolar epithelial cells.

*ADAM19* expression has also been shown to significantly decrease upon *PAX5* knock-out in B cells ^43^. Finally, another study has shown that hypomethylation of the *PAX6* motif located in the intronic region of *ADAM19* can increase *ADAM19* expression ^44^. Since the structure and docking of Pax5 is highly similar to Pax6 ^45^, it is possible that methylation of this site may impact Pax5 binding as well; however this would require further validation.

Previous work has suggested a crucial role for miRNA regulation in the differentiation, activation, and function of lymphocytes ^10,46^. Together with our findings of *PAX5* as main hub for miRNA regulation in blood, this led us to investigate miRNA-mRNA regulation in the context of B-cells. We leveraged B-cell specific deconvoluted expression data from the same cohort and repeated our network analysis pipeline, building subject-specific miRNA-mRNA regulatory networks and identifying interactions statistically associated with airflow obstruction. This analysis once again identified a very modular network, with two groups of miRNAs and two groups of mRNAs. The miRNA groups were similar to those identified in the bulk analysis, but there was little overlap in the mRNA groups. However, *PAX5* remained one of the top targeted mRNAs and we once again observed a strong differential expression signal, with mRNAs in each of the two groups collectively displaying either higher or lower expression as a function of decreasing airflow obstruction. In addition, gene set enrichment analysis on mRNAs differentially correlated with *PAX5* in the context of FEV_1_/FVC identified functions related to B-cell activation and differentiation. This, on the one hand, confirms the master regulatory role of *PAX5*. However, on the other hand, it points to a potential key role for *PAX5* in the innate immune response in COPD.

Finally, we analyzed a B-cell receptor (BCR) sequencing dataset which was previously collected on 137 subjects from COPDGene. We found statistical associations between many B-cell activation related functions and the expression of the network-identified genes, including *PAX5*. In particular, we found a significant (FDR < 0.05) positive association between *PAX5* expression and IgD and IgM counts, and a negative association between *PAX5* expression and IgA1, IgG1, and IgG2 counts, as well as with immunoglobulin class switching. Together, these findings confirms the key role of *PAX5* in pro-B and pre-B cells ^47^, specifically for IgM and IgD, and suggests a dysregulation of such responses in the context of COPD.

Limitations of our study include the sample source, which is whole blood rather than lung tissue, where the strongest inflammatory response against inhaled pathogens takes place. Future analysis should include testing in lung tissue in order to determine the role of *PAX5* in lung B cells. However, it is worth noting that blood is key for B-cell tissue infiltration, and we believe that the results presented in this manuscript likely reflect this process. Furthermore, these results may help to identify blood biomarkers for a dysregulated immune response in the case of pro-B and pre-B cells for COPD patients.

It is also worth noting that in our primary analysis we did not adjust for cell composition. However, when we conducted a sensitivity analysis correcting for white blood cell count and lymphocyte percentage, we still identified *PAX5* to be by far the most targeted gene. We also identified high targeting of *PAX5* in B cell specific deconvoluted expression data. As the data become available, it will be important to additionally leverage single cell RNA-seq to detail the relationship between *PAX5*, airflow limitation, and COPD in B cells. In addition, when evaluating *PAX5* targeting in the context of GWAS, we used the lead variants at genetic loci; for only some of these (e.g. *ADAM19*) we have reasonable confidence for the causal SNP. Finally, although our bulk and B cell specific deconvoluted data contained mRNA expression information for nearly 4000 subjects, only a small subset of these individuals (382) had miRNA expression and B cell receptor (137) data. This small sample size is clearly a limitation in our analyses that leverage miRNA expression information and may contribute to not obtaining clear differential expression signals for the miRNAs.

Overall, the results presented in this paper provide important insights into how miRNA-mRNA regulation in blood is impacted by airflow obstruction. In particular, we identified a key potential role for *PAX5* in mediating miRNA regulation of the innate immune response and B-cell activation in COPD. Furthermore, we found evidence of a direct regulatory interaction between *PAX5* and *ADAM19*, a COPD GWAS gene, and observed that this interaction may be impacted in the context of COPD. Together these findings point to potential mechanisms by which miRNAs, together with *PAX5*, regulate the innate immune response in COPD, and pave the way for future biomarker and therapeutic discovery.

## Acknowledgements

MG and KRG were supported by the National Institutes of Health (NIH), National Heart Lung and Blood Institute (NHLBI), through R01HL155749. MDM is supported by NIH/NHLBI K25HL168157. BDH was previously supported by NIH/NHLBI R01HL162813, R01HL155749, R01HL160008, U01HL089856, and a Research Grant from the Alpha-1 Foundation. CPH is supported by NIH/NHLBI R01HL166231. MLK is funded by the Norwegian Research Council, Helse Sør-Øst and the University of Oslo through the Centre for Molecular Medicine Norway (grant no. 187615), the Norwegian Research Council (grant no. 313932), the Norwegian Cancer Society (grant no. 214871 and 273592), and the iCAN Flagship in Digital Precision Cancer Medicine. MHC is supported by NIH/NHLBI R01HL160008, R01HL162813, and R01HL153248. MM is supported by NIH/NHLBI K08HL159318. PJC is supported by NIH/NHLBI R01HL124233, R01HL171213, R01HL166992.

## Code Availability

The python code of PUMA and LIONESS used to construct the miRNA-mRNA networks is available through netZooPy (https://github.com/netZoo/netZooPy). Additional code, including that used to analyze the networks, is available on GitHub (https://github.com/kimberlyglass/puma_copdgene).

