## Supplemental Materials, Methods, and Figures for "miRNA-mRNA network analysis identifies PAX5 as a potential regulator of adaptive immune response in COPD"

### Supplemental Materials and Methods

#### Additional Information on Data Processing

##### COPDGene RNA-seq and miRNA-seq expression data

Processing of whole blood RNA sequencing (RNA-seq) data (n = 3,985) from the Phase 2 (5-yr follow-up visit) of the COPDGene Study has been previously described in Ryu et al 2023[1]. Briefly, for samples passing quality control, isoform-level expression quantification was generated with Salmon (v1.3.0) pseudoaligned to GRCh38 and summarized to gene-level TPM (transcripts per million) counts using tximeta (v1.8.5), for a total count of 19,263 genes. TPMs were scaled to obtain gene length corrected counts and upper-quartile normalization was applied to obtain log-CPM (count per million) expression values that were further adjusted for library batch effects (removeBatchEffect, limma v3.46.0 and edgeR v3.32.1). For this analysis, we subsequently removed batches with less than 10 subjects, PRISm subjects, and never smoking subjects, for a final count of 3,190 subjects. We also only kept "protein coding" annotated genes (gencode v37) and filtered to only retain genes with an associated target mRNA in our TargetScan prior (see below). This resulted in a final data set containing 3,190 subjects and 11,859 genes.

As described previously[2] whole blood miRNA sequencing (miRNA-seq) data (n = 538) from Phase 2 of the COPDGene Study has been collected using single multiplexed blocking oligonucleotides were used to reduce unwanted hemolysis-related miRNAs hsa-miR-486-5p, hsa-miR-451a, and hsa-miR-92a-3p, with trimmed reads aligned to the hg19/miRBase 21 database. For this analysis, we performed additional preprocessing of these data similar to that performed in Zhuang et. al.[3]. Namely, we removed samples belonging to "plate5", with low sequencing depth (total read counts <200 k), or associated with subjects for which $FEV_1$/FVC was not defined, for a final count of 439 subjects. We also removed low expressed miRNAs by requiring more than 10 reads in at least 200 of these 439 subjects and a minimum standard deviation of 10 across subjects, for a final count of 679 miRNAs. We computed the trimmed mean of M-values (TMM) [4] and counts per million (CPM) normalization for these data using the *rnanorm* python package v 1.5.1.

##### miRNA target prediction data

To construct an input motif prior for the PUMA algorithm, we downloaded miRNA target predictions from TargetScan[5] v8.0 (all predictions, file "Summary Counts.all predictions.txt.zip," https://www.targetscan.org/vert_80/vert_80_data_download/Summary_Counts.all_predictions.txt.zip, accessed: July 17, 2023) and selected *Homo sapiens* interactions, resulting in a miRNA-mRNA network that included 2,606 miRNAs, 19,324 target mRNAs, and 10,011,946 edges. We then filtered to only include miRNAs and mRNAs (genes) expressed in the whole blood miRNA-seq and RNA-seq data, respectively (see above), resulting in a network that included 670 miRNAs, 11,859 mRNAs, and 1,779,028 edges. Each of these edges has an associated "TargetScan score", based on the sequence similarity (8mer) of the miRNA with a seed region on the target mRNA. miRNAs with high sequence similarity (≤ 2 base pair difference) belong to the same miRNA family. Therefore, to avoid high redundancy in our network (*i.e.* sets of miRNAs with nearly identical mRNA targets) we downloaded the definition of miRNA families from TargetScan 8.0 (https://www.targetscan.org/vert_80/vert_80_data_download/miR_Family_Info.txt.zip, accessed: July 17, 2023) and merged miRNAs belonging to the same family (see **Supplemental Table 1**) into a single node. If two miRNAs belonging to the same family had an edge to the same mRNA, we kept only the one with the lowest score (highest confidence). This resulted in a network composed of 570 miRNAs, 11,859 mRNAs, and 1,496,256 edges. We weighted the edges in this network by -1 times the TargetScan score, since the PUMA algorithm treats more positive values with higher confidence. We refer to this network as the TargetScan prior.

##### B-cell receptor sequencing data

Using bulk RNA-seq, we previously generated adaptive immune receptor repertoire sequencing data for B cell receptors (hereafter, 'BCR-seq') data using a set of isotype-specific immunoglobulin heavy chain (IGH) constant region primers[6]. Briefly, BCR-seq involves analyzing transcripts of the BCR by utilizing primers aimed at the Fc region of the immunoglobulin heavy chain (IGH). This method allows for a detailed evaluation of the BCR repertoire, covering antibody isotypes (IgM, IgD, IgA, IgG, and IgE), the variable region V-segments that define antibody specificity, as well as clonal expansion and somatic hypermutation within specific B cell groups. The data obtained is summarized into several quantitative indicators: 1) isotype usage (the proportion of antibody transcripts for each isotype per individual), 2) isotype expression (log2-transformed counts indicating the number of unique B cells per isotype per individual), 3) class switching (the percentage of class-switched B cells per individual), 4) V-segment usage (proportion of antibody transcripts for each V-segment per individual), and 5) CDR3 length by isotype. BCR-seq data has been collected from 240 subjects within the COPDGene Study [6]; 137 subjects with BCR-seq data also have bulk RNA-seq data (see above).

#### Additional Analysis: Network Robustness

To ensure that the subnetworks extracted in our analysis were not the result of a methodological artifact, we verified that upon randomization of the $FEV_1$/FVC phenotype we would not obtain any significant result. To test this, we randomized subject labels before running linear regression to identify edges statistically associated with $FEV_1$/FVC. We obtained only 2 statistically significant miRNA-mRNA interactions. This result illustrates that the observed subnetwork structure we obtain is due to combining information in the Target Scan prior, gene co-expression, and subject phenotype (i.e. $FEV_1$/FVC).

#### Supplemental Figures

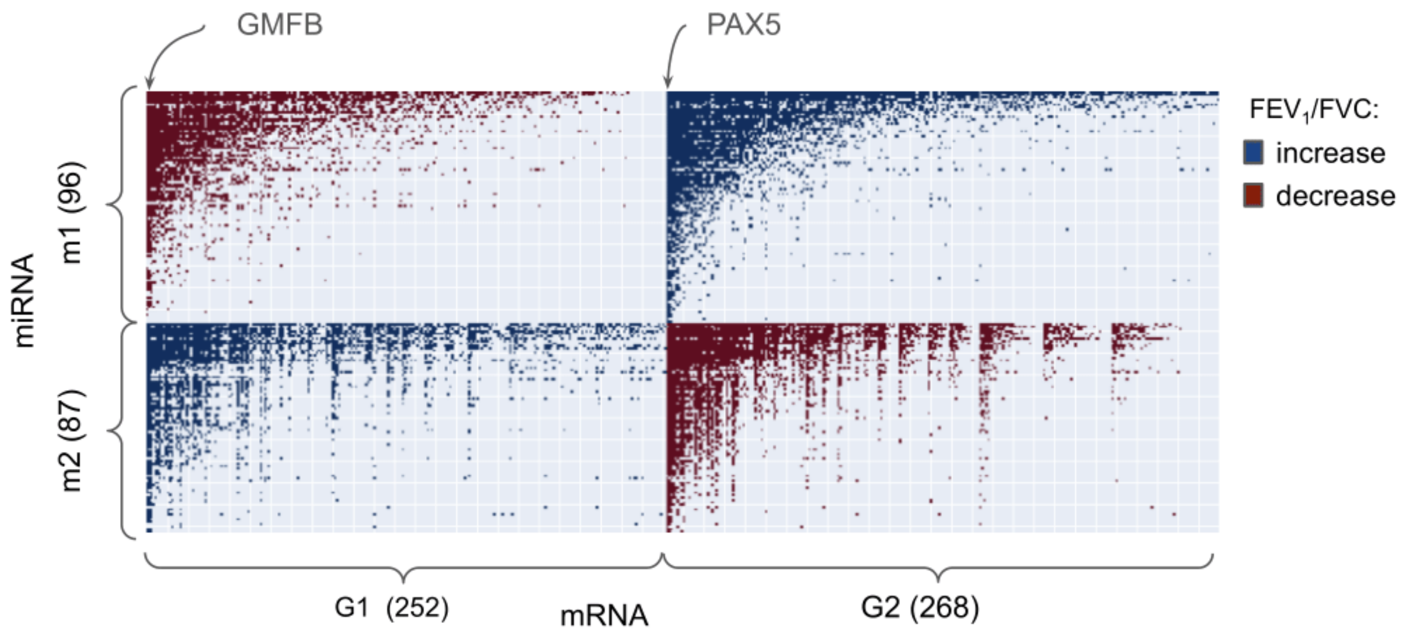


**Supplemental Figure 1: Network analysis using bulk whole blood expression data without filtering based on the TargetScan prior.** The clustered adjacency matrix of all the miRNA-mRNA interactions significantly associated with FEV$_1$/FVC. A dot in the matrix represents a significant edge between the miRNA (row) and mRNA (column). The color of the dot represents the sign of the association. Rows and columns have been sorted according to their degree and their subnetwork grouping.

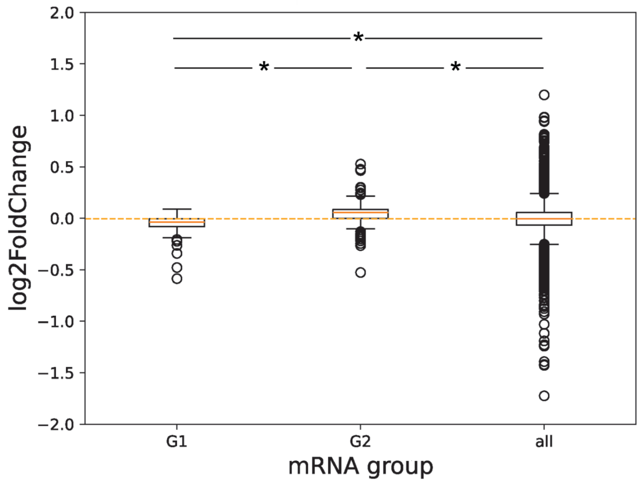


**Supplemental Figure 2**: **mRNA log2 fold change distribution**. The plot shows the distribution of the log2fold change computed for each mRNA using whole blood expression. We adjusted for age, sex, race, BMI, smoking status and cell composition (white blood cell count and lymphocyte percentage). An asterisk (*) indicates $p<0.05$ by two-sided t-test.

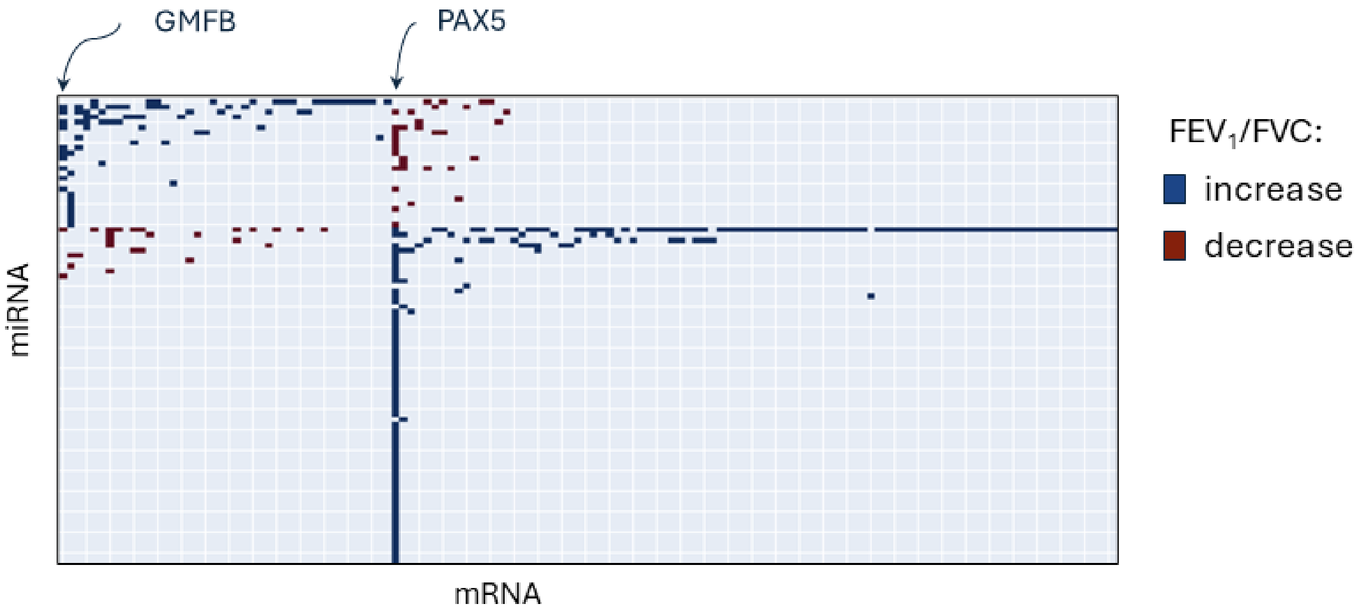


**Supplemental Figure 3:** The clustered adjacency matrix of all the miRNA-mRNA interactions significantly associated with $FEV_1$/FVC when additionally adjusting for cell composition (white blood cell counts and lymphocyte percentage). A dot in the matrix represents a significant edge between the miRNA (row) and mRNA (column). The color of the dot represents the sign of the association. Rows and columns have been sorted according to their degree and their subnetwork grouping.
